# Differential mitochondrial bioenergetics and cellular resilience in astrocytes, hepatocytes, and fibroblasts from aging baboons

**DOI:** 10.1101/2024.02.06.579010

**Authors:** Daniel A. Adekunbi, Hillary F. Huber, Cun Li, Peter W. Nathanielsz, Laura A. Cox, Adam B. Salmon

## Abstract

Biological resilience, broadly defined as ability to recover from acute challenge and return to homeostasis, is of growing importance to the biology of aging. At the cellular level, there is variability across tissue types in resilience and these differences likely to contribute to tissue aging rate disparities. However, there are challenges in addressing these cell-type differences at regional, tissue and subject level. To address this question, we established primary cells from aged male and female baboons between 13.3-17.8 years spanning across different tissues, tissue regions, and cell types including: (1) fibroblasts from skin and from heart separated into left ventricle (LV), right ventricle (RV), left atrium (LA) and right atrium (RA), (2) astrocytes from the prefrontal cortex and hippocampus and (3) hepatocytes. Primary cells were characterized by their cell surface markers and their cellular respiration assessed with Seahorse XFe96. Cellular resilience was assessed by modifying a live-cell imaging approach we previously reported that monitors proliferation of dividing cells following response and recovery to oxidative (50µM-H_2_O_2_), metabolic (1mM-glucose) and proteostasis (0.1µM-thapsigargin) stress. We noted significant differences even among similar cell types that are dependent on tissue source and the diversity in cellular response is stressor specific. For example, astrocytes were more energetic and exhibited greater resilience to oxidative stress (OS) than both fibroblasts and hepatocytes. RV and RA fibroblasts were less resilient to OS compared with LV and LA respectively. Skin fibroblasts were less impacted by proteostasis stress compared to astrocytes and cardiac fibroblasts. Future studies will test the functional relationship of these outcomes to age and developmental status of donors as potential predictive markers.

## 1.0 Introduction

Healthy cells undergo functional decline over time, with variations in the extent or timing of these changes attributed to their distinct morphological and physiological properties [1, 2]. Similarly, tissues and organs age at different rates and this cellular diversity is likely to contribute to these outcomes [3, 4]. In response to cellular perturbations, the homeostatic network orchestrates a coordinated dynamic response to restore stability. Cellular resilience measures the ability of a cell to recover to homeostasis following a challenge and can be used as functional readout of the homeostatic process. Central to the maintenance of cellular homeostasis is the pivotal role played by mitochondria, which provide the energy buffer required for appropriate stress responses and recovery through oxidative phosphorylation. A stabilized ATP supply enhances survival from stress, as well as the repair of damaged molecules to restore homeostasis [5]. Additionally, enhanced mitochondrial antioxidant capacity increases resilience to metabolic stress [6]. While energy demand and antioxidant defense capacity vary across different cell types, limited information exists concerning cellular resilience to stressors in different cell types.

Cell heterogeneity also increases with aging [7]. However, it is unclear whether these changes modify resilience or drive different resilience patterns across distinct cell types in an individual donor. That is, are cellular resilience properties conserved across the organ systems? This is important because understanding cell-specificity in homeostatic response may provide insights into basis of differences in tissue and organ aging rates. One study demonstrated that neonatal rat primary cardiomyocytes are more vulnerable to oxidative damage than cardiac fibroblasts, exhibiting marked DNA fragmentation and reduced cell survival following exposure to hydrogen peroxide [8]. Leveraging an ongoing aging study in a clinically relevant nonhuman primate (NHP) model [9,10], we here developed primary cell lines from aging baboons to address potential cellular resilience differences across multiple cell types from key organs of interest such as the brain, heart, and liver. The close genetic and physiological similarities of baboons to humans [11] enhances the potential applicability of our findings to humans.

In the present study, we assessed and compared the pattern of resilience across cell types in distinct organ regions and similar cell types within different anatomical locations in baboons. First, we derived and characterized resident cells from multiple regions of the heart (cardiac fibroblasts), brain (astrocytes), and liver (hepatocytes) of adult male and female baboons. We assessed mitochondrial response to its electron transport chain modulators using a mitochondrial stress test and assessment of bioenergetic function. We also incorporated a secondary test of resilience to mitochondrial stress test by exposing cells to low glucose to induce metabolic stress prior to the mitochondrial stress test. We also determined cellular proliferation response following short-term exposure to oxidative, proteostasis, and metabolic stress in the different cell types under the same stress intensity. This study lays the foundation for future investigations that will address the impact of age, sex, and developmental events on cellular resilience as well as other mechanistic studies within our baboon cohort.

## 2.0 Methods

### 2.1. Animals

All procedures involving animals were approved by the Institutional Animal Care and Use Committee of Texas Biomedical Research Institute (TX Biomed). The animal facilities at the Southwest National Primate Research Center (SNPRC), located within TX Biomed, are fully accredited by the Association for Assessment and Accreditation of Laboratory Animal Care International (AAALAC), which adheres to the guidelines set forth by the National Institutes of Health (NIH) and the U.S. Department of Agriculture.

Baboons (*Papio* species) were housed and maintained in outdoor social groups and fed *ad libitum* with normal monkey diet (Purina Monkey Diet 5LEO; 2.98 kcal/g). The welfare of the animals was enhanced by providing enrichment, such as toys, food treats, and music, which were offered daily under the supervision of the veterinary and behavioral staff at SNPRC. All baboons received veterinarian physical examinations at least twice per year and were healthy at the time of study.

The baboons were part of a cohort of animals studied throughout the life course to identify and establish effects of physiological aging [10,12–13]. All sample collections for primary cell cultures were taken opportunistically at prescribed euthanasia for animals in this larger study.

### 2.2. Sample Collection

Healthy male and female adult baboons aged between 13.3 and 17.8 years (approximate human equivalent, 50-70 years) were tranquilized with ketamine hydrochloride (10 mg/kg intramuscular injection) after an overnight fast, and then anesthetized with 1-2% isoflurane. While under general anesthesia baboons were euthanized by exsanguination as approved by the American Veterinary Medical Association. We have shown that euthanasia by intravenous agents can alter tissue structure [14]. Following cardiac asystole, all tissues were collected according to a standardized protocol within <1 hour from time of death, between 8:00-10:00 AM to minimize potential variation from circadian rhythms. Tissue collection was carried out under aseptic sterile conditions for isolation of resident primary cells. Skin biopsy was taken from behind the upper back portion of one ear of each baboon for isolation of epidermal fibroblasts. The prefrontal cortex was dissected from the left side of the brain. About 3 mm deep samples of gray mater from the prefrontal cortex, extending from the posterior part of the precentral sulcus to the intersection of the precentral sulcus and lateral sulcus were collected. Area 10 of the Broca’s area of the prefrontal cortex was processed to obtain cortical astrocytes. The whole hippocampus was carefully dissected from the temporal lobe and cut coronally into three pieces comprising anterior, middle, and posterior hippocampus, with about 2-3 mm of each piece combined for isolation of primary astrocytes. Cardiac tissues were collected separately from the left ventricle (LV), right ventricle (RV), left atrium (LA) and right atrium (RA) for cardiac fibroblast isolation. Both left and right liver lobes were collected separately for hepatocyte dispersion. All samples were collected on ice in sterile cell culture solution and processed within 1 h of collection.

### 2.3. Astrocyte cultures

The protocol for isolating baboon astrocytes was adapted from previous published procedures [15, 16]. Broca’s area 10 of the prefrontal cortex and hippocampus were minced in prewarmed 0.25% trypsin-EDTA (Gibco, Carlsbad, CA, USA) into small fragments. After 20 min of trypsinization in a standard cell culture incubator, an equal volume of DMEM-F12 Ham media (Sigma, St. Louis, MO, USA) supplemented with 10 % FBS and 1% antibacterial and antimycotic solution was added to the tissue fragments. The tissue was mechanically disrupted by passing through a serological pipette and pelleted by centrifugation at 1000x *g* for 2 min. Tissue fragments were further sheered by repeated pipetting. The processed mixture was then filtered through a 70 μm cell-strainer, and the filtered cells were plated in a tissue culture flask (surface area of 75 cm^2^) in a fully supplemented DMEM-F12 Ham media.

### 2.4. Fibroblast cultures

Skin and cardiac fibroblasts were obtained using a similar cell culture isolation procedure. We previously published a detailed protocol for isolating primary fibroblasts from baboon skin samples [9] which was adapted for isolating cardiac fibroblasts. Briefly, tissues were minced and incubated with liberase (0.4mg/ml, Sigma, St Louis, MO, USA) in a standard cell culture incubator for 3 h. Tissue mixture were resuspended in culture media, filtered with 100 µm cell strainer and centrifuge at 1000x *g* for 5 min. Dispersed cells were cultured in Dulbecco’s modified eagle medium, DMEM (Gibco, Carlsbad, CA, USA) supplemented with 10% fetal bovine serum (FBS, Gibco) and 1% antibiotic and antimycotic solution.

### 2.5. Hepatocyte cultures

The two-step EGTA/collagenase perfusion technique [17, 18] was adapted to isolate primary hepatocytes from baboon liver ex situ. Liver lobes were first separated from the entire liver piece and cut laterally closer to the caudal portion of the liver. Subsequently, two cannulae (16 g x 4 in) with a 3 mm smooth olive-shaped tip were positioned to target vascular channels in the liver for perfusion. We utilized a perfusion solution consisting of 1x HBSS-HEPES (comprising 0.14 M NaCl, 50 mM KCL, 0.33 mM Na_2_HPO4, 0.44 mM KH_2_PO4, 10 mM Na-HEPES) buffered with 0.5 mM EGTA, 5 mM Glucose, and 4 mM NaHCO_3_ and adjusted to pH 7.2 (EGTA solution). The perfusion was facilitated using a Masterflex peristaltic pump (Cole-Palmer, Niles, IL, USA) set at a rate of 10 revolutions per minute (rpm), and this procedure continued for approximately 1 hour at 37°C. The EGTA solution was devoid of calcium, as the presence of calcium prevents dissociation of the extracellular matrix for effective isolation of live hepatocytes.

Following perfusion with the EGTA buffer, the solution was replaced with a collagenase solution which comprises 0.1% collagenase, 5 mM CaCl_2_, and 4 mM NaHCO_3_ in 1X HBSS solution, pH 7.5. The liver was then perfused at a rate of 8 rpm for a duration ranging from 45 minutes to 1 hour depending on the liver size. The collagenase solution was collected and reused until visible signs of digestion were evident in the liver. A fully digested liver could be identified by its responsiveness to gentle pressure, resulting in an indentation, and the underlying cells are visible through the Glisson’s capsule that overlays the liver. The digested liver was collected into a 10 mm culture dish containing chilled Gibco’s Williams media supplemented with 5% FBS, 1% glutamine, and antibiotics. The liver cells were dispersed into the ice-cold media under a sterile cell culture hood by gently removing the Glisson’s capsule and teasing apart the tissues with a 1 ml pipette tip. Cell suspensions were filtered through sterile folded gauze, centrifuged at 50 *g*, 4°C for 5 min. The centrifugation step was repeated twice, and the resulting hepatocytes were plated on a collagen-coated plate containing prewarmed media overnight before performing further experiments the next day.

### 2.6. Characterization of primary cells with cell surface markers

Immunocytochemistry was used to validate cell type identity. Cells were fixed using 4% v/v paraformaldehyde, then rinsed with 1X HBSS and permeabilized with 0.1% Triton X-100. Cells were blocked with 5% bovine serum albumin in TBS with 0.1% Tween20 at room temperature for 1 h and then incubated in primary antibodies (detailed in Table 1) overnight at 4 °C. These primary antibodies were selectively targeted towards markers associated with fibroblasts (vimentin), hepatocytes (cytokeratin 8) and astrocytes (aquaporin 4, glial fibrillary acidic protein (GFAP), and excitatory amino acid transporter 2; EAAT2). Markers of neurons (post synaptic density protein 95; PSD95), and oligodendrocytes (myelin basic protein) were used to confirm whether isolated astrocyte population are mixed with other brain cell types. Cultures were probed with secondary antibodies conjugated with FITC, counterstained using DAPI for nuclear identification, and imaged with a Zeiss LSM 880 confocal microscope using a 63x oil immersion lens.

**Table 1:** Primary antibodies specific for cell surface markers of fibroblasts, hepatocytes, and astrocytes.

| Cell Marker | Antibody | Dilution ratio | Company | Cat # |
| --- | --- | --- | --- | --- |
| Fibroblasts | Vimentin | 1:200 | Santa Cruz | sc-6260 |
| Hepatocytes | Cytokeratin 8 | 1:200 | Santa Cruz | sc-8020 |
| Astrocytes | aquaporin 4 | 1:200 | Millipore | AB2218 |
|  | GFAP | 1:200 | Millipore | MAB360 |
|  | EAAT2 | 1:200 | Santa Cruz | sc-365634 |
| Neurons | PSD95 | 1:200 | Millipore | MABN1190 |
| Oligodendrocytes | MBP | 1:100 | Cell Signaling | 78896 |

### 2.7. Seahorse mitochondrial assay

Agilent Seahorse XFe96 Extracellular Flux Analyzer (North Billerica, MA, USA) was used to measure cellular respiration. Cells were plated in a 96 well seahorse plate at density of 40,000 cells per well for all the different cell types. For hepatocytes requiring an adhesion material for cell attachment to the plate, collagen derived from rat tail (Sigma) was prepared in sterile double distilled H_2_O in 1:50 ratio and incubated in seahorse plates at 37 °C for 3 h before cell seeding. The XFe96 sensor cartridges were hydrated overnight with H_2_O at 37 °C and replaced with XF calibrant 1 h before the assay. Oxygen consumption rate (OCR), an index of cellular respiration was measured under basal condition and in response to serial injection of mitochondrial inhibitors including 1.5 µM Oligomycin (to inhibit ATP synthase), 0.5 µM FCCP (Carbonyl cyanide-p-trifluoromethoxyphenylhydrazone; a mitochondrial uncoupler to measure maximum respiration) and 0.5 µM antimycin A/rotenone (to inhibit electron flow through mitochondrial electron transport chain). To determine mitochondrial response to metabolic stress, a subset of cells was exposed to a low glucose media (1 mM glucose) for 2 h, before the mitochondrial stress test described above. After the assay, the OCR values were normalized to cell density measured by IncuCyte live-cell imaging system. Data were analyzed by the Agilent WAVE software.

### 2.8. Cellular resilience assay

Cellular resilience was assessed in dividing primary cells (except hepatocytes which do not proliferate in vitro). Cell proliferation rate was determined in response to 50 µM hydrogen peroxide (H_2_O_2_), low glucose media (1 mM), or 0.1 µM thapsigargin to model oxidative, metabolic and proteostasis stress respectively using a live-cell imaging system (IncuCyte S3). Cells were plated at a density of 2,000 cells per well into a 96-well plate with cell growth captured every 6 h for 6 d. After 24 h of cell plating, cells were exposed to H_2_O_2_, low glucose and thapsigargin for 2 h. Treatments were replaced with complete media and growth monitored until the termination of the study. The slope of the time-course changes in cell confluence particularly during the exponential growth phase following exposure to stressful challenges was used to determine cell proliferation rate (% confluence per h) using GraphPad prism 9 software.

### 2.9. Statistical analysis

Data were analyzed by two-way analysis of variance (ANOVA) followed by Tukey post hoc test and presented as mean ± SEM; *p* < 0.05 is considered statistically significant. Data from males and females were combined, n=5-10 baboons per tissue cell type except where they are specifically separated by sex. The variability in animal number per tissue type was due to failed culture of some cell types from a single donor. All analyses were carried out using GraphPad prism 9.

## 3.0 Results

### 3.1. Characterization of baboon primary astrocytes, fibroblasts, and hepatocytes

Astrocytes derived from the prefrontal cortex and hippocampus show strong immunoreactivity for astrocyte markers; aquaporin 4, glial fibrillary acidic protein (GFAP), and excitatory amino acid transporter 2 (EAAT2). The absence of immunoreactive signals for myelin basic protein and post synaptic density protein 95 (PSD95) indicates that astrocyte cultures do not contain oligodendrocytes and neurons, respectively. Both skin and cardiac fibroblasts showed strong immunoreactive signal to vimentin, a marker of fibroblasts. Phase contrast image showed hepatocytes as polygonal cells with distinct cell border. The hepatocytes were immunopositive for cytokeratin 8 (Fig. 1).

**Fig. 1:**
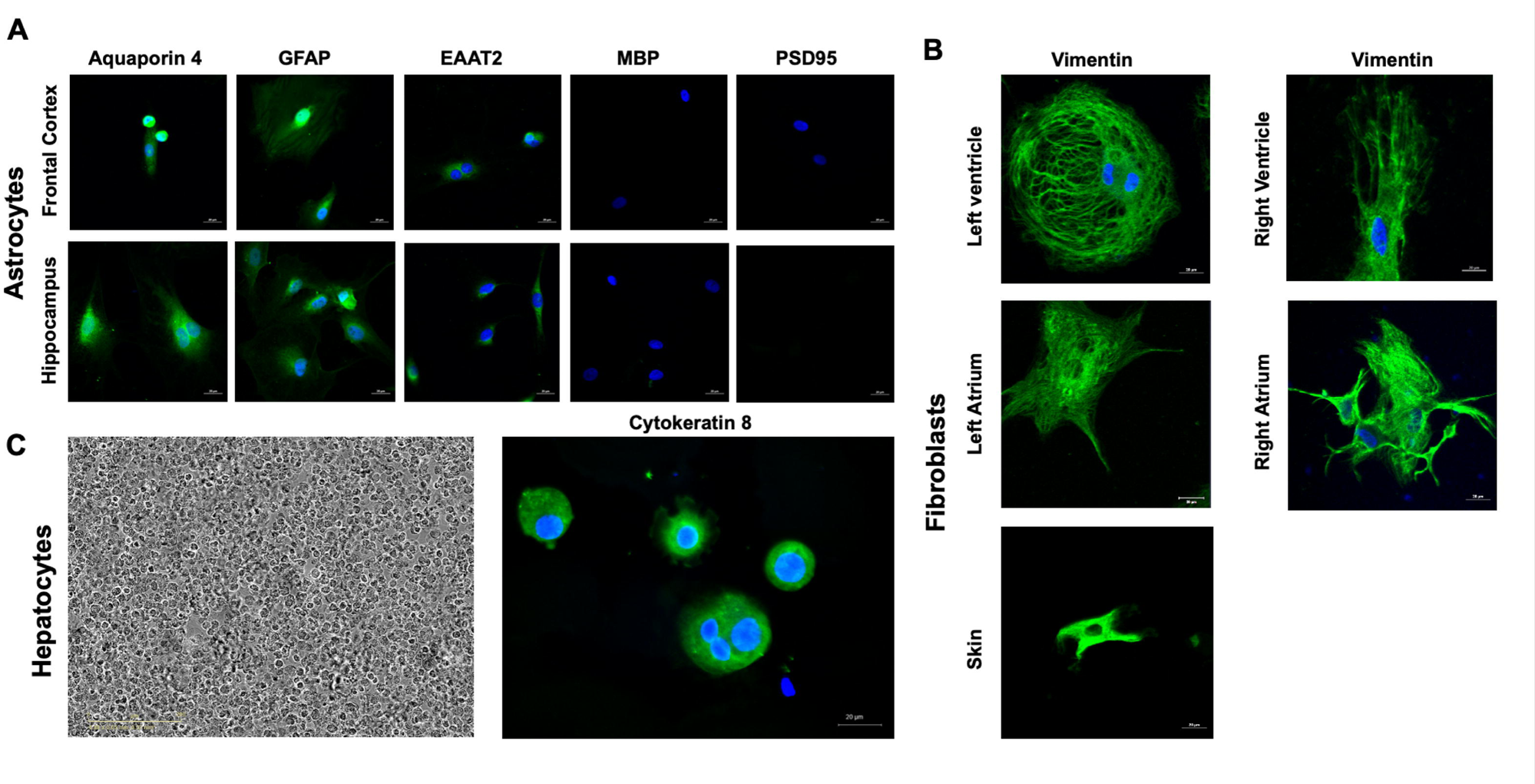
Representative photomicrographs of baboon astrocyte, fibroblast, and hepatocyte cultures. (A) Astrocytes derived from prefrontal cortex and hippocampus show immunoreactivity for astrocyte markers; aquaporin 4, glial fibrillary acidic protein (GFAP), and excitatory amino acid transporter 2 (EAAT2). The absence of immunoreactive signals for myelin basic protein and post synaptic density protein 95 (PSD95) indicates that astrocyte cultures do not contain oligodendrocytes and neurons, respectively. (B) Expression of fibroblast marker, vimentin in fibroblasts from different heart regions such as the left ventricle, right ventricle, left atrium, and right atrium as well as skin epidermal fibroblasts express fibroblast marker, vimentin. (C) Phase contrast microscopy of hepatocyte cultures and immunoreactive signals of epithelial cell marker, cytokeratin 8. Scale 20 µm. Images acquired with a Zeiss LSM 880 confocal microscope using a 63x oil immersion lens. Donor animal: Male, 16.4 years old.

### 3.2. Mitochondrial bioenergetics differ among astrocytes, fibroblasts, and hepatocytes

In figure 2 (A-E), we present OCR kinetics in astrocytes, fibroblasts, and hepatocytes under normal culture conditions (designated as control) and in response to acute metabolic stress induced by exposure to low glucose (1 mM glucose for 2 h). These were combined data for male and female. With the exception of astrocytes, the OCR of other cell types examined was comparable under both control culture conditions and those under metabolic stress prior to testing. However, by testing all under the same conditions we show the differential respiratory patterns, both basal and under mitochondrial stress during the assay, among the different cell types from aging baboon cells. Figure 2 F shows the cellular respiration pattern of these 5 different cell types under control conditions. Further, figures 2 G-I show the integrated data and significant differences among these cell types in their basal, ATP-linked, and maximal respiration properties.

**Fig. 2:**
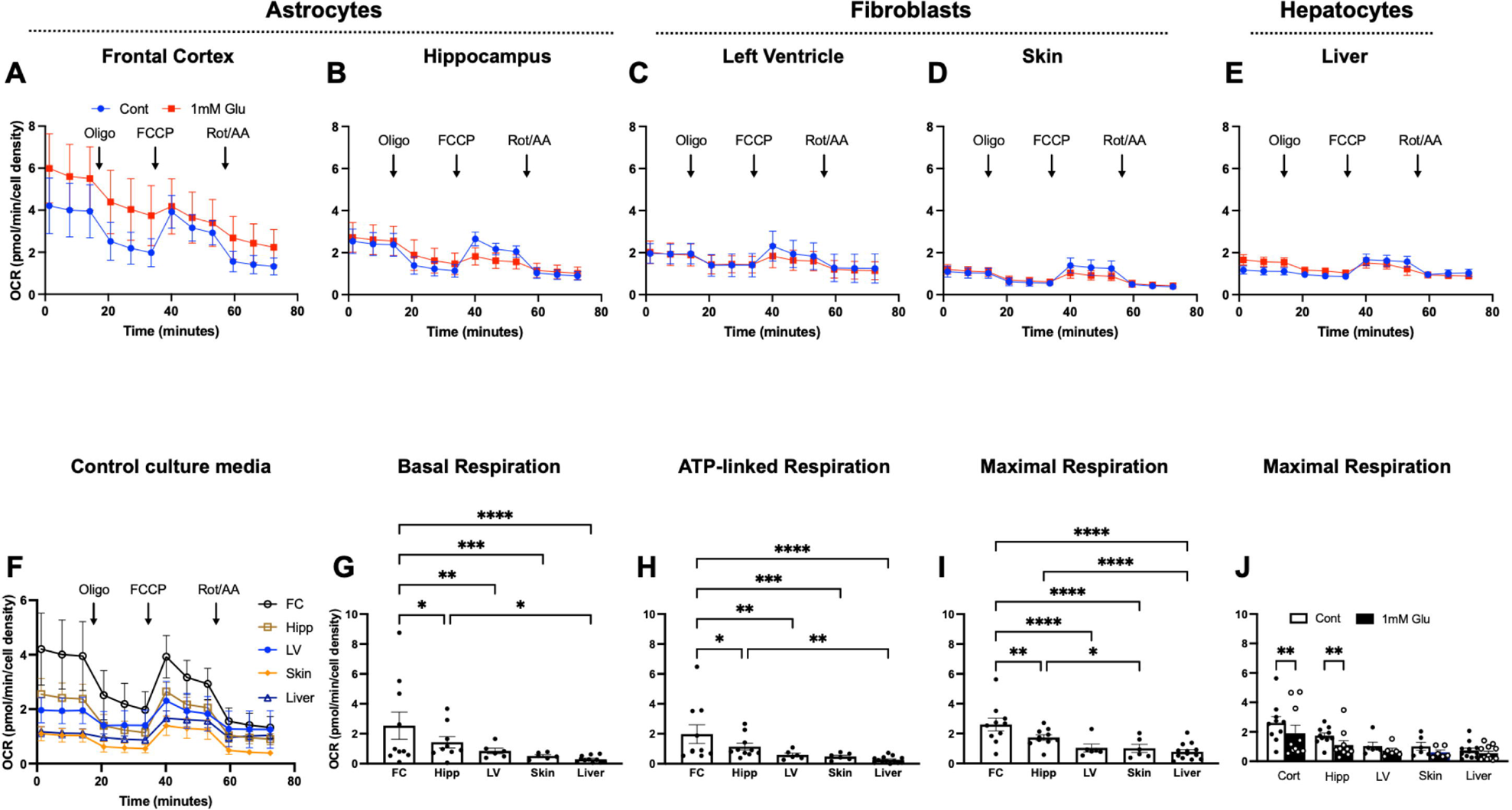
Oxygen consumption rate in baboon astrocytes, fibroblasts, and hepatocytes. The kinetics of oxygen consumption rates (OCR) in (A) astrocytes derived from the Broca’s area of the prefrontal cortex (FC), (B) astrocytes from the hippocampus, (C) fibroblasts from the left ventricle of the heart, (D) skin fibroblasts and (E) hepatocytes under standard culture condition and in response to low glucose (1mM glucose for 2h). (F) Time-course OCR of cortical astrocytes, hippocampal astrocytes, LV fibroblasts, skin fibroblasts, and hepatocytes under standard culture conditions. (G) Basal respiration, (H) ATP-linked respiration, and (I) Maximal respiration in astrocytes derived from the FC and hippocampus (hipp) as well as left ventricle fibroblasts (LV), skin fibroblasts (skin) and hepatocytes (liver). Data from individual animal were expressed as mean ± standard error of mean, with male and female data combined. Data from left and right lobe hepatocytes were pooled for comparison with other cell types. For each primary cell line, respiration rates were from 4 to 6 replicate samples measured using the seahorse XFe96 flux analyzer. Donor age, 13.3-17.8 years, sample size (cortical astrocytes, n=10; hippocampal astrocytes, n=10. LV and skin fibroblasts, n=6 respectively; hepatocytes, n=7. *p<0.05, **p<0.01, ***p<0.001, ****p<0.0001.

Cortical astrocytes exhibited the highest overall OCR among the cell types, followed by hippocampal astrocytes, while OCR in LV and skin fibroblasts as well as hepatocytes were comparable. There was a significantly higher basal, ATP-linked, and maximal respiration in cortical astrocytes relative to hippocampal astrocytes, LV fibroblasts, skin fibroblasts, and hepatocytes. Hippocampal astrocytes exhibited significantly higher bioenergetic profiles compared to skin fibroblasts and hepatocytes but not with LV fibroblasts (Fig. 2 G-H). Interestingly, astrocytes were sensitive to acute exposure to low glucose, evidenced by reduced maximal respiration in both cortical and hippocampal astrocytes, whereas other cell types did not exhibit changes.

While we found no effect of metabolic stress (low glucose) on respiratory outcomes in fibroblasts and hepatocytes, we did see the astrocytes from both cortex and hippocampus show reduced maximal respiration when cultured under metabolic challenge (Fig. 2 J). We used LV cardiac fibroblasts as representative of this tissue/cell type because we found no significant difference in mitochondrial respiration among cell lines isolated from the heart (discussed below). No significant difference in bioenergetics was observed between left lobe and right lobe hepatocytes (data not shown); thus, data from both lobes were pooled for comparison with other cell types.

We next asked whether donor sex affected these cellular and mitochondrial phenotypes. Data from both males and females were relatively consistent with the data analyzed as combined sexes above (Supplemental Fig. 1). We see the same cell-type specific respiration patterns across cell types when data are analyzed for only male and for only female donors. Comparison by sex showed that only female cortical astrocytes had higher maximal OCR compared to males while other cell types exhibited similar OCR between male and female donors (Supplemental Fig. 1 C).

### 3.3. Comparison of astrocyte and fibroblast resilience to oxidative, metabolic and proteostasis stress

Using a protocol we previously developed for epidermal derived fibroblasts [9], we measured cellular resilience in response to different stressors such as H_2_O_2_ (50 µM), low glucose (1 mM), and thapsigargin (0.1 µM) to model oxidative, metabolic and proteostasis stress respectively. For these studies, hepatocytes were excluded due to their lack of proliferation under standard culturing conditions. Baboon astrocytes and fibroblasts from different anatomical regions exhibited similar proliferation rate when cultured in standard culture media or exposed to low glucose media but they responded quite differently to oxidative or proteostasis stress (Fig. 3). The proliferation of astrocytes from both the prefrontal cortex and hippocampus was not altered by short-term exposure to 50 µM H_2_O_2_ whereas RV and skin fibroblasts had reduced proliferation in response to a 2h H_2_O_2_ challenge suggesting increased cellular resilience to challenge in astrocytes. Interestingly, the impact of oxidative stress on LV fibroblasts was similar to that of astrocytes. The exposure of astrocytes and fibroblasts from different anatomical region to low glucose media for 2 hours inhibited cell growth in a comparable pattern, resulting in no discernible difference between the different cell types. However, with thapsigargin challenge, the resulting reduction in proliferation rate was significantly greater in astrocytes, RV and LV fibroblasts compared to skin fibroblasts suggesting low resilience to proteostatic stress in these cell types (Fig. 3).

**Fig. 3:**
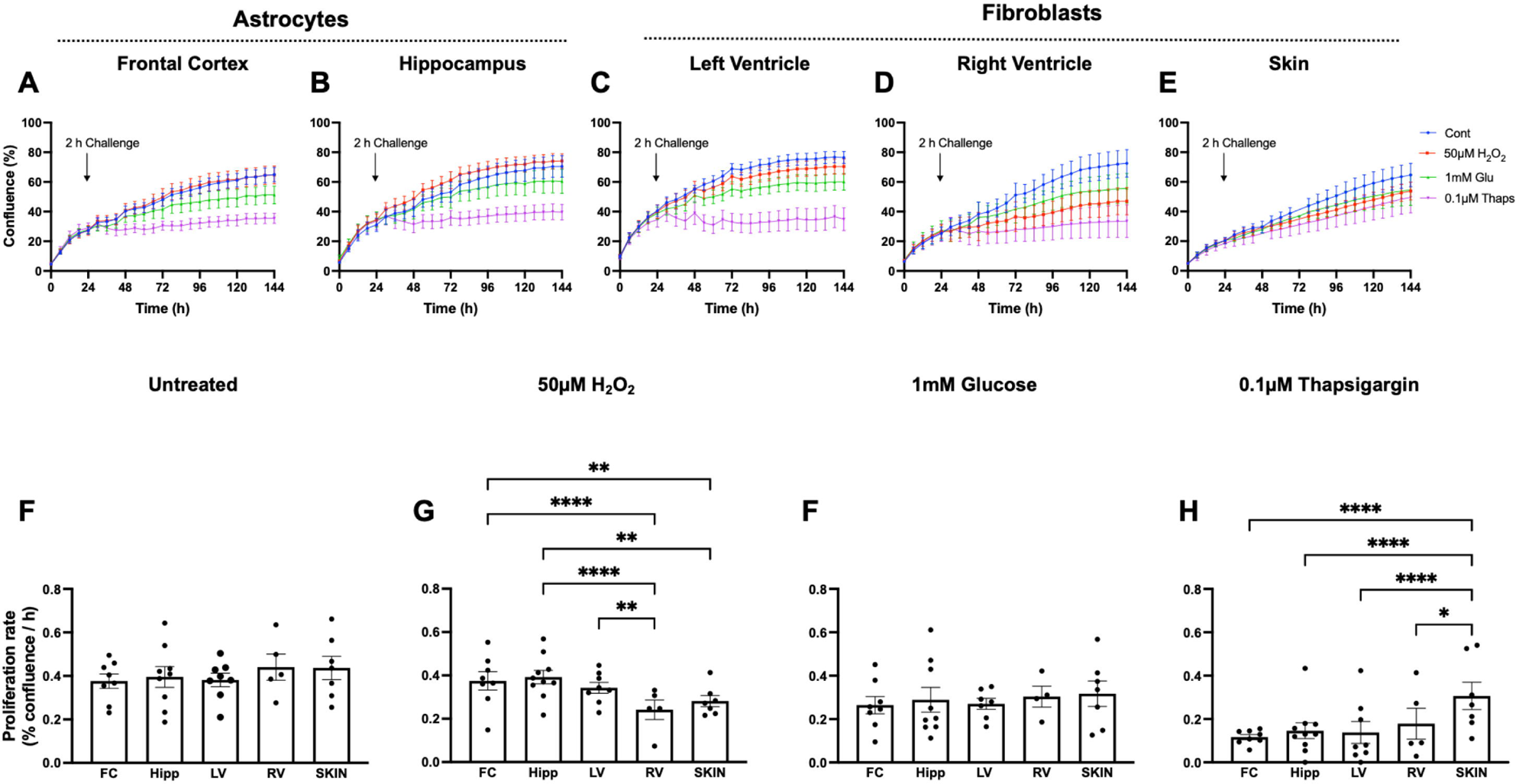
Effects of oxidative, metabolic and proteostasis stress on proliferation of baboon astrocytes and fibroblasts. Astrocytes derived from prefrontal cortex Broca’s area 10 (FC) or the entire hippocampus (hipp) encompassing anterior, middle, and posterior hippocampus and fibroblasts from the left ventricle (LV), right ventricle (RV) and ear skin (skin) were exposed to 50µM H_2_O_2_, 1mm glucose, and 0.1 µM thapsigargin for 2h to model oxidative, metabolic and proteostasis stress respectively. Time-course changes in cell confluence in response to challenge were monitored real time with the IncuCyte live-cell imaging system housed within a cell culture incubator (3% O_2_, 5% CO2 at 37 °C). The blue line represents untreated cells (designated as control), red line; 50µM H_2_O_2_, green; 1mM glucose and purple, 0.1µM thapsigargin. Proliferation rate (% confluence/h) calculated from the slope of the kinetic graph between 30 and 144 h were shown in response to untreated and challenged cells. Proliferation rate was analyzed using two-way ANOVA. Data expressed as mean ± standard error of mean, each data point represents 3 replicate wells for each animal, cell seeding density; 2000 cells/well, donor age were between 13.3 and 17.8 years, with male and female data combined. Sample size: FC, n=8; hipp, n=10; LV, n=8; RV, n=5; skin, n=7. *p<0.05, **p<0.01, ****p<0.0001.

Similar to our mitochondria assays above, we found only small effects of donor sex on these outcomes in these data sets. The response of the different cell types to stressors were similar in male and female subjects except the slight variation in the response of cortical astrocytes and RV fibroblasts to oxidative and proteostasis stress respectively between males and females (Supplemental Fig. 2).

Given the higher metabolism of astrocytes and their resilience to 50µM H_2_O_2_ challenge, we reasoned that there may be a relationship between bioenergetics and resilience of primary astrocytes. Indeed, astrocyte maximal respiration significantly correlates with their proliferation response to H_2_O_2_ challenge (Fig. 4), which suggests higher maximal respiration may be predictive of astrocyte resilience to oxidative stress.

**Fig. 4.**
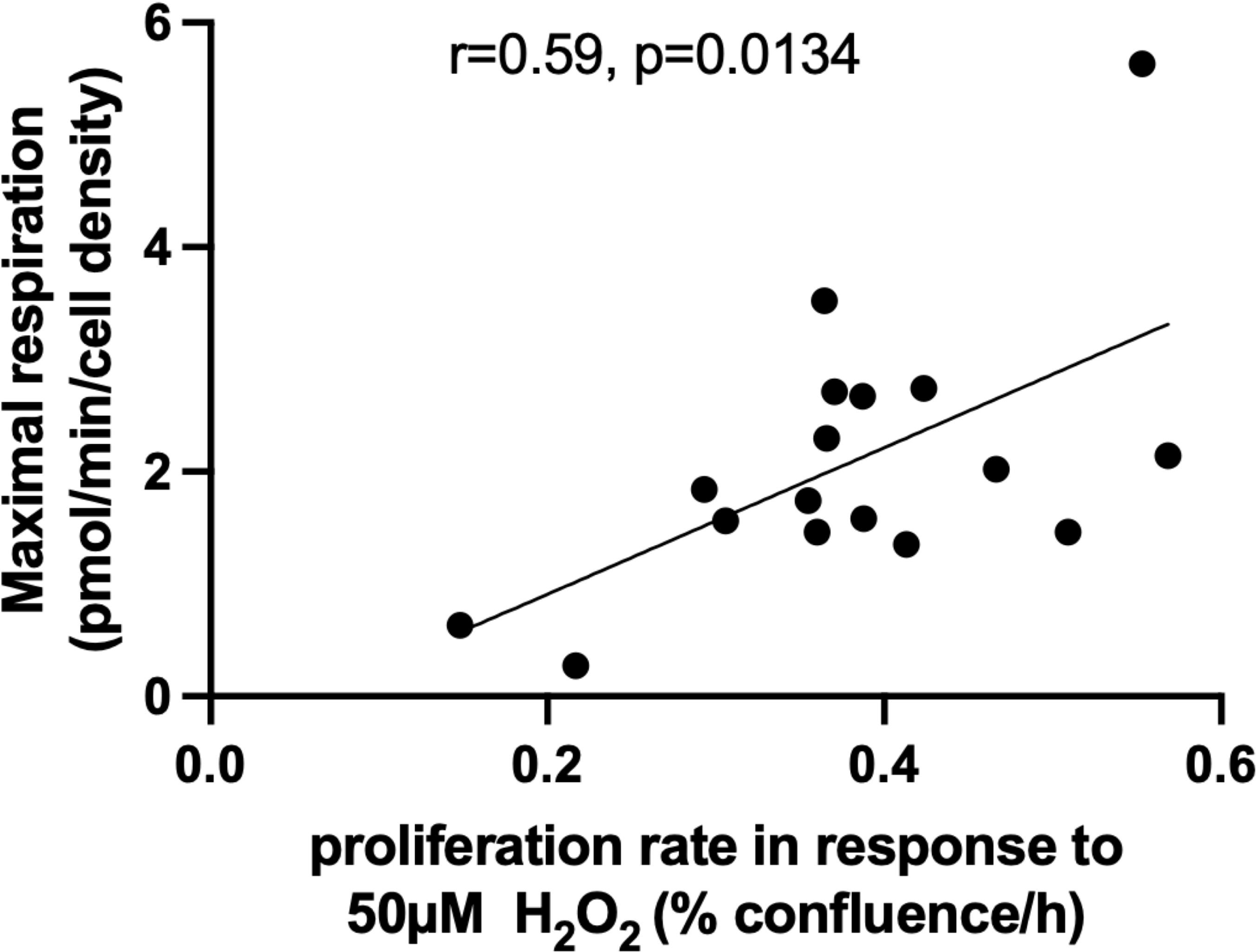
Correlation between astrocyte maximal respiration and proliferation response to oxidative stress. The maximal oxygen consumption rates (OCR) in baboon astrocytes, induced by the mitochondrial uncoupler FCCP, shows a positive correlation with their proliferation response to 50µM H_2_O_2_. OCR was determined with the seahorse XFe96 flux analyzer. Cellular proliferation was monitored with a live-cell imaging system (IncuCyte) with the proliferation rate calculated from the slope of cell kinetics over 5 d period. The correlation between maximal respiration and proliferation rate in response to H_2_O_2_ was determined using Pearson’s correlation coefficient. Data are from primary astrocytes obtained from baboon prefrontal cortex and hippocampus encompassing both male and female animals, with ages ranging from 13.3 to 17.8 years (n=17).

### 3.4. Mitochondrial bioenergetics in cardiac fibroblasts from different heart regions

Even within a single tissue, cells are not homogenous in cell type, distribution, location, or niche. Here we addressed whether cells isolated from different regions of the same tissue (heart) might differ in patterns of resilience. We made comparison between primary fibroblasts from heart chambers to determine whether site-specific differences exist in cell bioenergetics. Cardiac fibroblasts from different heart regions (LV, RV, LA, and RA) exhibited similar OCR for all measures of seahorse mitochondrial stress test under standard cell culture conditions (25 mM glucose in culture media) or following 2 h exposure to low glucose media to model acute metabolic stress. Figure 5 (A-D) shows OCR kinetics in LV, RV, LA, and RA fibroblasts from which basal, ATP-dependent, and maximal respiration were derived (Fig. 5 E-F). These data then suggest, at least in the heart, that anatomical location has little impact on respiratory function of primary cells.

**Fig. 5:**
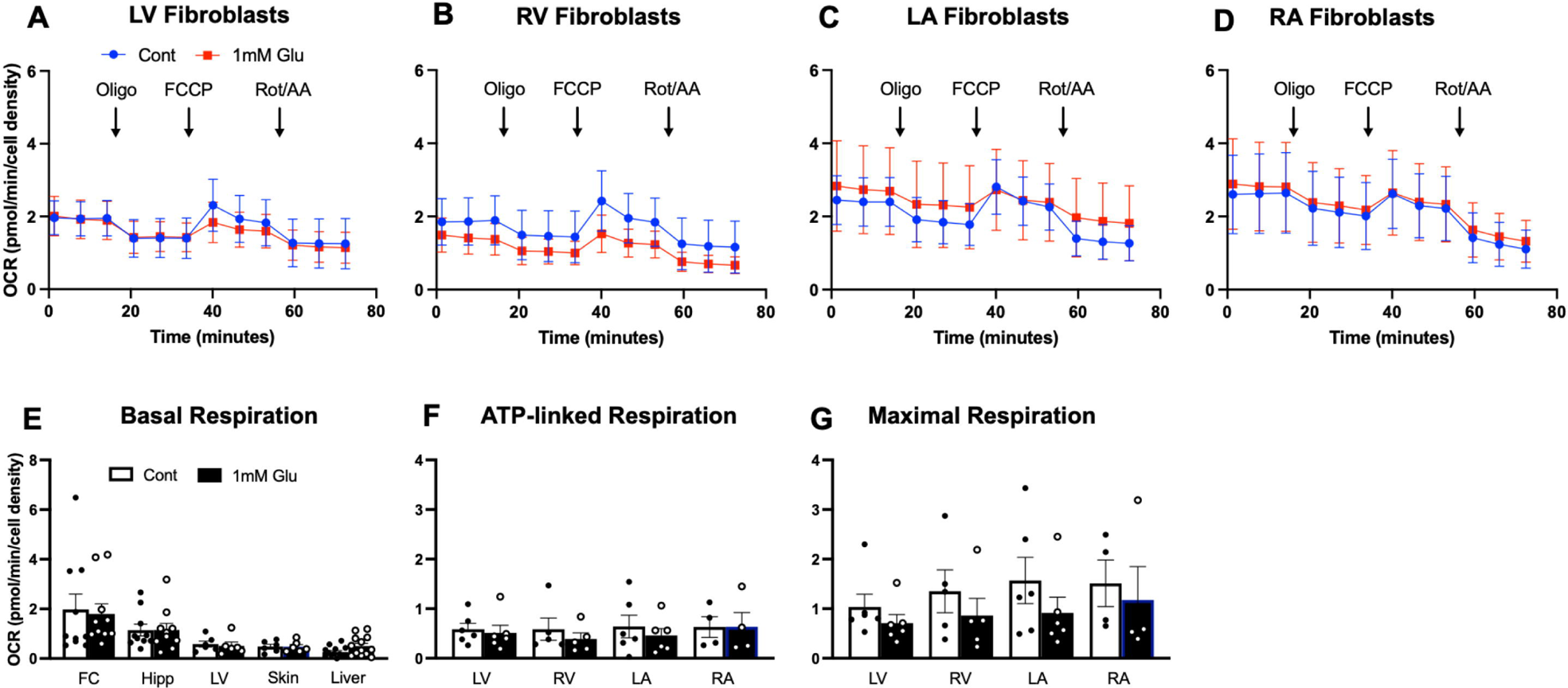
Oxygen consumption rate of baboon cardiac fibroblasts. Time-course graph of oxygen consumption rates (OCR) measured with a seahorse XFe96 flux analyzer using baboon cardiac fibroblasts derived from (A) left ventricle; LV, (B) right ventricle; RV, (C) left atrium; LA and (D) right atrium; RA, under standard culture condition (designated as control) and in response to low glucose media (1mM glucose). (E) Basal respiration, (F) ATP-linked respiration and (G) Maximal respiration were derived from the kinetic graph following initial measurement, in the presence of oligomycin and in response to Carbonyl cyanide-p-trifluoromethoxyphenylhydrazone (FCCP) respectively. Data, expressed as mean ± standard error of mean, are from individual animal, with male and female data combined. For each primary cell line, respiration rates from 4 to 6 replicate samples were measured using the seahorse XFe96 flux analyzer. Donor age, 13.3-17 years, n=5-6 per cardiac fibroblasts from different heart regions.

### 3.5. Resilience of cardiac fibroblasts to oxidative, metabolic and proteostasis stress

On the other hand, we did find evidence that resilience to challenge among cells within a tissue may be impacted by anatomical location of development. Figure 6 shows the kinetics and proliferation rate of cardiac fibroblast in normal growth media and in response to challenge compounds. Cardiac fibroblasts from different heart regions exhibited similar proliferation rates under normal culture conditions (Fig. 6 A-E). However, upon short-term exposure to H_2_O_2_, the proliferation of both RV and RA fibroblasts reduced significantly compared to the effect of H_2_O_2_ on LV and LA fibroblasts, suggesting that fibroblasts from the right heart region are less resilient to oxidative stress (Fig. 6 A-D and F). When subjected to either metabolic or proteostasis stress, (1 mM glucose or 0.1 µM thapsigargin, respectively) cardiac fibroblasts demonstrated impaired proliferation in a similar manner across different heart regions (Fig. 6 A-D, G and H).

**Fig. 6:**
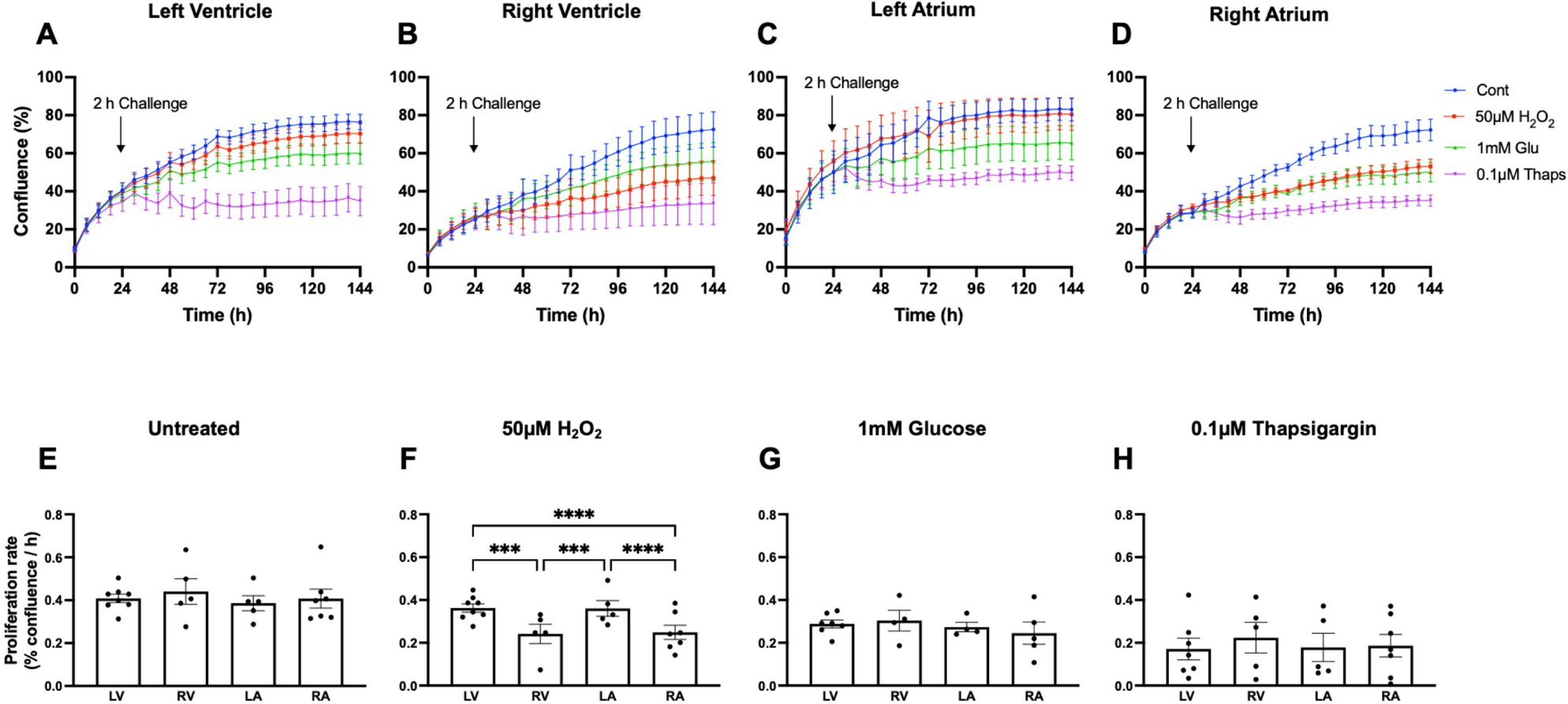
Effects of oxidative, metabolic and proteostasis stress on proliferation of cardiac fibroblasts from different heart region. Fibroblasts derived from the four chambers of baboon heart encompassing left ventricle (LV), right ventricle (RV), left atrium (LA) and right atrium (RA) were exposed to 50µM H_2_O_2_, 1mm glucose, and 0.1 µM thapsigargin for 2h to model oxidative, metabolic and proteostasis stress respectively. Time-course changes in cell confluence of cardiac fibroblasts in response to challenge compounds were monitored real time with the IncuCyte live-cell imaging system housed within a cell culture incubator (3% O_2_, 5% CO2 at 37 °C). The blue line represents untreated cells (designated as control), red line; 50µM H_2_O_2_, green; 1mM glucose and purple, 0.1µM thapsigargin. Proliferation rate (% confluence/h) calculated from the slope of the kinetic graph between 30 and 144 h were shown in response to untreated and challenged cells. Proliferation rate was analyzed using two-way ANOVA. Data expressed as mean ± standard error of mean, each data point represents 3 replicate wells for each animal, cell seeding density was 2000 cells/well, donor age between 13.3 and 17.8 years, male and female data combined, n=8 for LV, 5 for RV, 5 for LA and 7 for RA. ***p<0.001, ****p<0.0001.

## 4.0 Discussion

The newly established cell lines described in this study represent a valuable resource for mechanistic studies that can be coupled with in vivo approaches to bridge some translational gaps between basic and clinical studies in aging research and other allied fields. This work is part of an ongoing initiative to generate different primary cell types from an aging baboon cohort across the life-course. The data presented herein pertain specifically to adult baboons aged between 13.3 and 17.8 years and thus do not ask specifically whether age changes these patterns of resilience; primary cells from younger baboons, including various cell types beyond skin fibroblasts will be tested in a similar manner in the near future. We characterized the identity of multiple tissue cultures in baboons and determined their bioenergetic and resilient properties under similar experimental conditions. In most instances, the sex of the donor animal did not significantly alter the observed differences between cell types. The contribution of male donors complements that of females and the combination of male and female added more statistical power. As a result, we premised data interpretation primarily on combined data from both sexes, which aligns with the results obtained from individual sex-based analyses for most variables. Significant findings here include the following: (i) Prefrontal cortex astrocytes exhibit higher mitochondrial respiration than hippocampal astrocytes. (ii) Astrocytes are more energetic and exhibit greater resilience to oxidative stress than both fibroblasts and hepatocytes. (iii) Metabolic stress alters maximal respiration only in astrocytes but changes in fibroblasts and hepatocytes were not statistically significant. (iv) Fibroblasts obtained from the right heart region (RV and RA) were less resilient to oxidative stress compared to fibroblasts from the left heart region (LV and LA). (v) Skin fibroblasts were less impacted by proteostasis stress relative to astrocytes and cardiac fibroblasts. (vi) Higher maximal respiration may serve as a predictive indicator of astrocyte resilience to oxidative stress.

We characterized the identity of primary cells with their cell surface markers using immunocytochemistry. Astrocyte cultures exhibited strong reactivity to key astrocyte markers such as GFAP, aquaporin 4, and EAAT2, and importantly, they did not contain mixed populations of oligodendrocytes and neurons. Similarly, both skin and cardiac fibroblasts demonstrated robust immunoreactivity for vimentin, a recognized marker of fibroblasts, while hepatocytes were immunopositive to cytokeratin 8, a marker for epithelial cells. It is worth noting that cardiac fibroblasts are larger in size and morphologically different than skin fibroblasts (Fig 1). Whether these differences translate to any functional difference is not clear. The choice of these cell types is premised on other parallel studies on developmental programming and aging interactions in our baboon cohort where we observed accelerated aging of the brain, cardiac remodeling, and metabolic dysfunction due to developmental exposure to maternal undernutrition [19-21). Although neuronal and cardiomyocyte cultures are desirable, there are technical limitations of generating viable neurons or cardiomyocytes in aging NHP. Beside fibroblast cultures, there are few studies that have established primary cultures of key organs from baboons [22] and to our knowledge this study is the first to establish primary cells from multiple regions of the heart, brain, and liver in aging baboons (13.3-17.8 years). Baboon lifespan at SNPRC, where our baboons are maintained, has been reported as 11 or 21 years [23, 24] and thus these animals represent the transition period from mid- to late-life. This additional unique baboon resource has the potential to contribute to the development of translatable strategies to promote healthy aging and mitigating age-related diseases.

As a key determinant of cellular health, the mitochondria play a significant role in energy metabolism, metabolite production as well as cellular signaling [25]. However, their contribution to various cellular processes varies across different cell types, depending on the cell’s specific demands, differences in mitochondrial protein composition, and internal mitochondrial structures such as matrix and cristae [26]. In heart and brain cells, the mitochondria are predominantly oriented towards energy metabolism, converting substrates such as glucose to ATP to sustain high energy demanding tasks of the heart and brain. The liver in contrast plays a more significant role in metabolite biosynthesis and its mitochondria have lower metabolic dynamic range for energy conversion compared to those of the heart and brain. Mitochondria with a relatively higher abundance of cristae, where the majority of electron transport chain (ETC) complexes and ATP synthase dimers are located, possess greater capacities for energy conversion. The heart has more cristae compared to the liver, whose mitochondria prioritize a larger matrix volume over cristae to favor greater biosynthesis and metabolite signaling [26, 27].

Using cellular OCR as a proxy for mitochondrial respiration, we observed similar mitochondrial respiration levels between cardiac fibroblasts and hepatocytes. Given that myocardial contraction, an energy-dependent process, is determined by cardiomyocytes, it is plausible that the differences between heart and liver mitochondria may be closely related to the cardiomyocytes than cardiac fibroblasts. Astrocytes exhibited higher mitochondrial respiration than fibroblasts (both cardiac and skin) and hepatocytes. This finding may be attributed to the abundance of mitochondria in astrocytes and the overall high energy metabolism observed in the brain [27]. Although neurons consume about 80% of energy in the brain, most of which are expended on synaptic signaling, astrocytes play a significant role in ensuring energy delivery from capillaries to synapses, as such, astrocytes are considered as key cells in coupling synaptic activity with energy metabolism [28]. Given the role that astrocytes play in the energetic requirement of neurons, it is unsurprising that astrocytes exhibit higher mitochondrial respiration compared to other examined cell types. Furthermore, only astrocytes were sensitive to acute low glucose exposure, demonstrating greater glucose utilization and demand in astrocytes compared to fibroblasts and hepatocytes. This aligns with higher glucose metabolism in astrocytes compared to neurons [29] and parallels the essence for brain glycogen content storage mostly in astrocytes [30].

When we considered brain region specific variation in astrocytic mitochondrial function, cortical astrocytes exhibited higher basal, ATP-linked, and maximal respiration compared to hippocampal astrocytes. These functional disparities align with inherent morphological differences between the two astrocyte populations. In a morphometric analysis of differentiated cortical and hippocampal astrocytes derived from neural progenitor cells, cortical astrocytes displayed numerous short, well-defined projections with multiple branches per projection and smaller soma areas. In contrast, hippocampal astrocytes possess larger soma areas with few but long projections with almost no branching [31]. The distinctions in astrocytic projections, which is crucial for energy processing in support of neuronal metabolism through delivery of substrate from capillaries to synapse, may underlie the heightened energetic profile of cortical astrocytes compared to hippocampal astrocytes. Determining mitochondrial function in astrocytic branches will be important to demonstrate the bioenergetics of these astrocytic projections.

Cardiac fibroblasts play a crucial role as supporting cells for cardiomyocytes, analogous to the role of astrocytes in relation to neurons, although their specific supporting functions differ. As part of the myocardial wall, cardiac fibroblasts are involved in synthesis and maintenance of extracellular matrix (ECM) network which is essential for electrical conductivity and heartbeat rhythm [32]. We did not observe any region-specific differences in mitochondrial respiration of cardiac fibroblasts, which is consistent with other reports that have demonstrated that the processes of energy metabolism including mitochondrial protein contents in the left and right heart are identical [33, 34].

Interestingly, fibroblasts from the right side of the heart (RV and RA) were less resilient to H_2_O_2_ (inducer of oxidative stress) compared to those from the left region (LV and RA). It is not clear whether the nearly constant aeration of the left heart region with oxygenated blood influences stress response. We speculate that the region-specific response to oxidative stress may be related to stress adaptation in the left side of the heart, possibly influenced by the differential pressure loads between the heart chambers. For example, the LV works against higher pressure to pump oxygenated blood through the peripheral systemic vasculature while the RV works against low pressure to deliver deoxygenated blood to the lungs [35]. The elevated pressure load in the LV is a form of mechanical stress and pressure overload is linked to increased formation of superoxide anions [36]. Our results suggest that LV fibroblasts may have developed a reserve capacity for stress resilience compared to the RV. Indeed, others have reported that ROS-induced damage occur earlier in RV than LV due to lower antioxidant capacity in RV tissues [37]. Thus at the cellular level, the differential response to acute H_2_O_2_ challenge between the right and left heart region resonates with the intrinsic difference in their functional capacity. This differential response of cardiac fibroblasts from different heart regions is however specific to oxidative stress as there were no significant differences in proliferation response of cardiac fibroblasts to metabolic and proteostasis stress. We did not test this resilience assay in hepatocytes since they non-dividing cells *in vitro*.

When we compared the effect of H_2_O_2_ on proliferation of astrocyte and fibroblasts from different regions, H_2_O_2_ significantly decreased the proliferation of RV and skin fibroblasts compared to both cortical and hippocampal astrocytes. The effect of acute exposure to H_2_O_2_ was similar in astrocytes and LV fibroblasts. In contrast, skin fibroblasts were less impacted by thapsigargin exposure compared to astrocytes, LV, and RV fibroblasts. This suggests opposing sensitivity to oxidative and proteostasis stress in different cell types, an important consideration for future investigations aimed at between defining the molecular factors regulating resilience across tissue types. The relationship we observed between astrocyte bioenergetics and proliferation response to oxidative stress propels the idea that we can model interconnected hallmarks of resilience that would serve as predictive markers for clinical applications. The hallmarks may encompass variables such as mitochondrial bioenergetics, proteostasis, transcriptional response among others. In astrocytes, a higher maximal respiration is predictive of resilience to oxidative stress.

In summary, we provide evidence of successful establishment of astrocytes, fibroblasts, and hepatocyte cultures from our aging baboon cohort and that these cells display differential response to multiple forms of stress and mitochondrial bioenergetics. This cellular heterogeneity implies differential mechanisms underly stress resiliency across cell types which may impact their aging rates. A limitation of this study was the relatively small sample size in some cell types when examining the impact of donor sex on the variables studied. Our future studies will determine the role of sex in larger sample size, age, and developmental exposures on resilience across different cell types as we increase animal sampling.

## Acknowledgements

This research was funded in part by R01 AG057431, I01BX004167 (ABS), the San Antonio Nathan Shock Center (P30 AG 013319) and the Geriatric Research, Education and Clinical Center of the South Texas Veterans Health Care System. This material is the result of work supported with resources and the use of facilities at South Texas Veterans Health Care System, San Antonio, Texas. The contents do not represent the views of the U.S. Department of Veterans Affairs or the United States Government. Baboons in this study were maintained under 1U19AG057758-01A1 (PWN, LAC). The authors acknowledge the administrative and technical support of Karen Moore, Wenbo Qi, Benjamin Morr, and Rachel Camones. We also acknowledge support from the SNPRC which is funded by P51 OD011133.

## Competing Interest

The authors declare no competing interest.

## Notes

### Competing Interest Statement

The authors have declared no competing interest.

