## Supplemental Fig. 1, Supplemental Fig. 2 for "Differential mitochondrial bioenergetics and cellular resilience in astrocytes, hepatocytes, and fibroblasts from aging baboons"

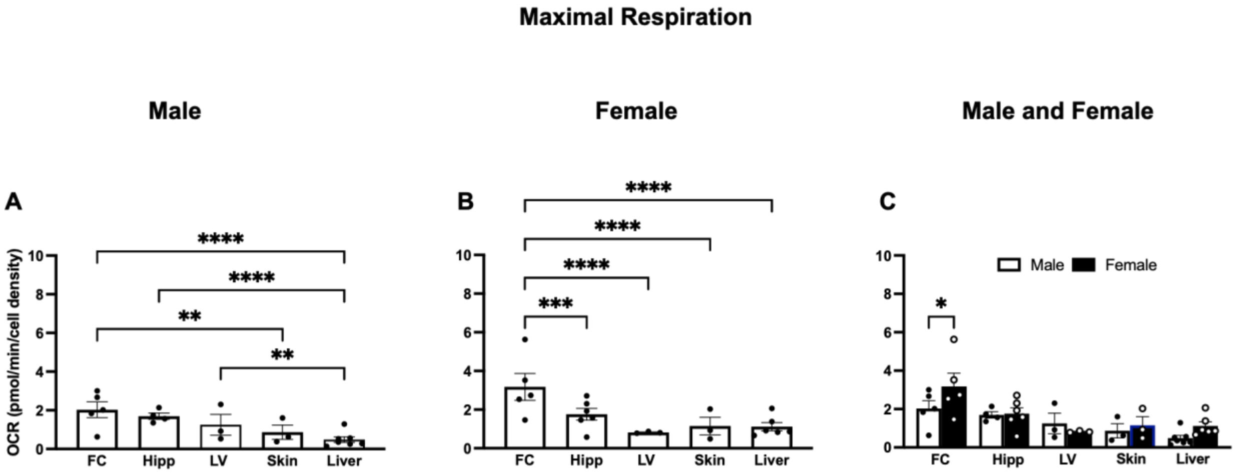


**Supplemental Fig. 1**: Sex differences in maximal respiration of baboon astrocytes, fibroblasts, and hepatocytes. Maximal oxygen consumption rates (OCR) were determined in astrocytes derived from the prefrontal cortex (FC) and hippocampus (hipp), left ventricle fibroblasts (LV), skin fibroblasts (skin) and hepatocytes (liver). (A) Maximal respiration in male baboons. (B) Maximal respiration in female baboons. (C) Comparison of maximal respiration between male and female baboons. Maximal respiration for each cell line were measured using the seahorse XFe96 flux analyzer in 4 to 6 replicate samples and expressed as mean ± standard error of mean. Donor age, 13.3-17.8 years, sample size (FC, n=5 each for males and females; hipp, n=4 for males, 6 for females; LV and skin fibroblasts, n=3 each for both sexes respectively; hepatocytes, n=4 for males, 3 for females. *p<0.05, **p<0.01, ***p<0.001, ****p<0.0001.

**Supplemental Fig. 2**

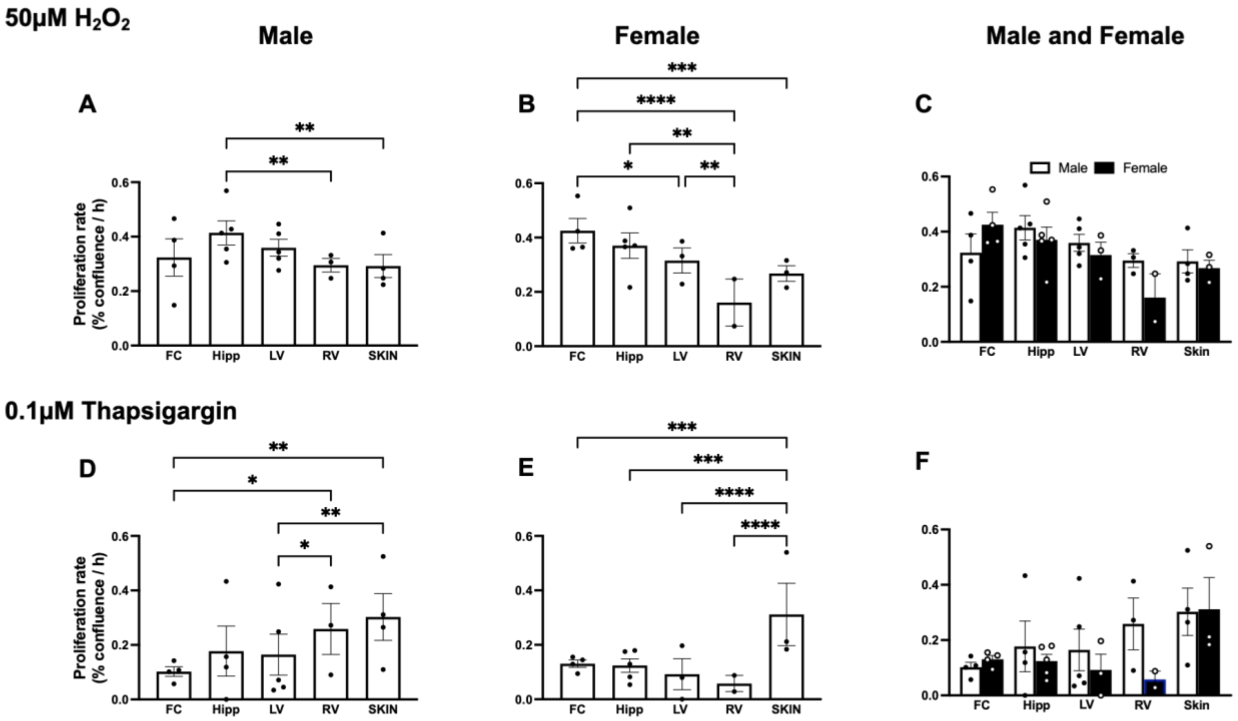


**Supplemental Fig. 2**: Comparison of sex differences between astrocyte and fibroblast responses to oxidative and proteostasis stress. Proliferation rate of astrocytes from the prefrontal cortex (FC) and hippocampus and fibroblasts from the left ventricle (LV), right ventricle (RV) and ear skin (skin) were determined following 2h exposure to $H_2O_2$ (50µM) or thapsigargin (0.1 µM) in male and female baboons. (A) Cellular proliferation rates in response to $H_2O_2$ challenge in males. (B) Cellular proliferation rates in response to $H_2O_2$ challenge in females. (C) Comparison of proliferation response to $H_2O_2$ challenge in male and female baboons. (D) Cellular proliferation rates in males following thapsigargin challenge. (E) Proliferation rates in response to thapsigargin challenge in females. (F) Comparison of proliferation response to thapsigargin challenge in male and female baboons. Data expressed as mean ± standard error of mean, donor age were between 13.3 and 17.8 years. Sample size: FC, n=4 per sex; hipp, n=5 per sex; LV, n=5 for males, 3 for females; RV, n=3 for males, 2 for females; skin, n=4 for males, 3 for females. *$p<0.05$, **$p<0.01$, ***$p<0.001$, ****$p<0.0001$.
